# LMPred: Predicting Antimicrobial Peptides Using Pre-Trained Language Models and Deep Learning

**DOI:** 10.1101/2021.11.03.467066

**Authors:** William Dee

## Abstract

Antimicrobial peptides (AMPs) are increasingly being used in the development of new therapeutic drugs, in areas such as cancer therapy and hypertension. Additionally, they are seen as an alternative to antibiotics due to the increasing occurrence of bacterial resistance. Wet-laboratory experimental identification, however, is both time consuming and costly, so in-silico models are now commonly used in order to screen new AMP candidates. This paper proposes a novel approach of creating model inputs; using pre-trained language models to produce contextualized embeddings representing the amino acids within each peptide sequence, before a convolutional neural network is then trained as the classifier. The optimal model was validated on two datasets, being one previously used in AMP prediction research, and an independent dataset, created by this paper. Predictive accuracies of 93.33% and 88.26% were achieved respectively, outperforming all previous state-of-the-art classification models.

## Introduction

Antimicrobial peptides (AMPs) are a set of naturally-occurring molecules that exhibit a wide range of functions, including antibacterial, anticancer, antifungal and antihypertensive properties (1). When used to create therapeutic drugs, peptides are increasingly showing efficacy in terms of treatment in a variety of important areas; ranging from cancer targeting, to eliminating bacterial, viral and fungal pathogens (2).

Current cancer therapies, such as radiotherapy and chemotherapy, often elicit harmful side-effects, and cancer cell resistance to chemotherapeutic agents is a large and growing issue (3). Peptides have long been used as biomarkers in the detection and diagnosis of specific cancers, such as pancreatic, colorectal and lung (4). However, recently their use has been extended, as they have been shown to bind to specific cancerous sites and so have been used as carriers for targeted drugs (5). Certain peptides have also been shown to exhibit an inhibitory effect in cancer cells themselves (6). AMPs with anticancer properties (ACPs) are therefore promising drug development candidates due to their “high specificity, low production cost, high tumor penetration, and ease of synthesis and modification” (7), as well as their low toxicity.

Additionally, pathogenic bacteria are more frequently developing multi-drug resistance (1, 8), rendering current antibiotic treatments ineffective. Given that AMPs are endogenous, they have shown a lower likelihood for bacteria to develop resistance to them (9), and so offer a complementary alternative to traditional drugs.

Identification of natural AMPs is therefore becoming increasingly important. However, experimental identification is both time-consuming and costly, hence the need for in-silico prediction models (1, 10). Previous model-based methods have utilized a range of Machine Learning approaches, including; Hidden Markov Models (11), Fuzzy K-Nearest Neighbour (FKNN) (4), Random Forest (RF) (1), Discriminant Analysis (DA) (12) and Support Vector Machines (SVM) (7, 13, 14). Whilst these methods achieved high levels of accuracy, the feature creation steps were often extensive. Bhadra et al. (1) performed a survey of existing approaches to summarize the common data preprocessing steps. Previous research was most likely to use “compositional, physicochemical, structural properties, sequence order and the pattern of terminal residues” in order to create the final feature set.

Manavalan et al. (13) generated 20 features representing amino acid compositon, 400 relating to dipeptide composition, 5 associated with atomic composition – i.e., the frequency of each atom (C, H, N, O, and S) within each peptide sequence – and 11 referring to physicochemical properties – i.e., the percentage composition of polar, hydrophobic, charged, aliphatic, aromatic, positively/negatively charged, tiny/small and large residues, as well as overall peptide mass. In total this resulted in 436 additional features.

To address the large, and often sparse, feature spaces produced by prior methods, Bhadra et al. used an approach based on the “Global Protein Sequence Descriptors” (15). This created a reduced set of 105 features which was subsequently cut to 23 using a correlation analysis based on feature importance measures. At the time this was considered the “smallest feature set for AMP prediction with high accuracy” (1).

Compositional metrics, like those described above, fail to take sequence order into account – even though it is integral to the underlying function of peptides. To account for this, some prior approaches used Chou’s pseudo-amino acid composition (PseAAC) (16). This generates correlation factors between each amino acid, to partly incorporate some information relating to sequence order (4, 17). Similarly, Evolutionary Feature Construction (EFC) (18) has also been used to detect functional signals representing nonlocal, position-specific interactions at the nucleotide level (19).

Two recent papers have achieved improved model accuracies by applying Natural Language Processing (NLP) techniques to create vectorized embeddings of the amino acid sequences as the sole input features (20, 21). The embeddings are fed into a convolutional neural network in order to classify sequences as functional AMPs or not. This has the benefit of reducing the time taken, as well as the expert biological knowledge needed, when creating model inputs. Furthermore, NLP approaches have the potential to represent richer positional information reflecting the specific locations of amino acids within a given sequence.

Veltri et al. (20) utilized the Bag of Words (BoW) method to initially assign a unique numerical token to each amino acid in a sequence. This approach can recognize basic patterns, such as which amino acids are the same, and the frequency of specific amino acids in a sequence or throughout the dataset. However, BoW fails to capture component similarity. To partially alleviate this issue, an initial embedding layer was used in the neural network architecture, to convert the discrete vectors into a continuous and dense latent space represented by a three number vector, which can then reflect more complex relationships between inputs. Wu et al. (21) used the Word2Vec Skip-Gram algorithmn which learns by being given a central word (or amino acid in this case) and then predicting the most likely words in a fixed window surrounding it, allowing it to reflect word similarity (22). However, this method still fails to convey the contextual information encoded by the position of each amino acid.

This paper proposes a novel method of creating embedding vectors - by utilizing language representation models that have been pre-trained on large protein databases to produce *contextualised* embeddings. Language Models (LMs) are built using the Transformer architecture (23), and have primarily been used for language-based tasks, given that they have been pre-trained on large corpora such as the 2,500 million words found in Wikipedia (24).They have been proven to outperform humans in some assessments, such as the Stanford Question Answering Dataset – a reading comprehension test (25), as well as the two NLP approaches previously mentioned. When these models are pre-trained on protein sequences, treating each amino acid as a word, and the sequence as a sentence, they have shown an ability to generalize towards understanding the “language of life” itself (26). The LMs selected for this report are the auto-encoder models BERT (24) and T5 (27), and the auto-regressive model XLNet (28). These have already been pre-trained using the Summit supercomputer on either the UniRef100 or the UniRef50 datasets, consisting of 45 and 216 million protein sequences respectively (29). The UniRef clusters are proteins sourced from the UniProt database, with the number referring to the similarity threshold set in the CD-HIT program (30). Therefore, UniRef100 contains all UniProt sequences, whereas UniRef50 only contains sequences that do not share >=50% identity. Additionally, BERT and T5 were also pre-trained on the Big Fat Database dataset, comprising 2,122 million sequences (31). All models were accessed via the ProtTrans Github page (https://github.com/agemagician/ProtTrans). Bidirectional Encoder Representations from Transformers (BERT) aims to predict data from artificially corrupted inputs. It does this by adding [MASK] tokens during pre-training to replace a percentage of input words at random and it then aims to predict this masked word based on the surrounding context. In this manner it is able to capture bidirectional context. However, all masked words are predicted in parallel and independently of one another, which means that some of the overall context of a sentence can be lost during training. Also, whilst the artificial [MASK] tokens are used during pre-training, they are absent when the model is subsequently fine-tuned on a specific task, which can result in a “pretrain-finetune discrepancy” (28).

XLNet is an auto-regressive model, which functions more similarly to a feed forward neural network; these models aim to predict the next word from a set of words, given the context. However, unlike auto-encoder models, the prediction is constrained to be uni-directional. XLNet partially overcomes this drawback through permutation language modelling, whereby the model maximizes the expected log likelihood of the sequence when all possible permutations of the sequence order are considered (28). By learning context from randomly ordered sentences the resulting model is in essence bidirectional without requiring masking. Furthermore, XLNet utilizes a memory mechanism introduced by a previous auto-regressive model - Transformer-XL (32) - which allows for processing of longer contextual segments of data than BERT.

The Text-To-Text Transfer Transformer (T5) was introduced after Raffel et al. (27) studied the current landscape of transfer learning techniques for NLP and found that, generally, encoder-decoder models outperformed those only utilizing one half of the Transformer architecture. The T5 model therefore uses both parts of the Transformer, in contrast to BERT which only uses the encoder, and XLNet which only uses the decoder. T5 was also trained on a new “Colossal Clean Crawled Corpus”, a comparatively *clean* and *natural* text dataset that was several magnitudes larger than many previous training datasets. There is evidence that this additional complexity, and the fact that positional encodings for each attention head can be shared across all layers, gives T5 a performance advantage over the other two LMs (33).

A convolutional neural network has been chosen as the classifier based on its success in prior peptide prediction research (20, 21), as well as its successful application to other areas within bioinformatics, such as mapping protein sequences to folds (34) or the prediction of RNA secondary structure (35). The convolutional layers apply filters which can interpret the spatial and temporal dependencies between amino acids that have been represented by the contextualized LM embeddings.

## Methods

### Datasets

Validation of the approach was performed on two datasets. One which was sourced externally and has been used in prior AMP prediction research - referred to as the “Veltri Dataset”. The second has been constructed independently using publicly available resources - the “LMPred Dataset”.

### Veltri Dataset

The Veltri dataset was extracted from the Antimicrobial Peptide Scanner web page (https://www.dveltri.com/ascan/). This contains 1,778 AMP and 1,778 non-AMP samples and is split into the exact training, validation and test sets that Veltri et al. (20) used to build and evaluate their model.

### LMPred Dataset

The LMPred dataset has been built from a combination of external sources. The aim was to produce the largest and most up-to-date AMP dataset possible.

The positive samples (the AMPs) have been gathered from the freely available datasets shared by Veltri et al. (20) and Bhadra et al. (1), and combined with the peer-reviewed, natural AMPs taken from the DRAMP 2.0 database (36). When duplicates were removed, these sources contributed 7,053 AMPs. The samples are then filtered by removing AMPs less than 10 amino acids in length, as well as those sharing 90% sequence identity according to the CD-HIT online web server (30). CD-HIT is a program used to cluster proteins according to a defined similarity criterion. Applying the program to the dataset can help to eliminate bias towards particular arrangements of sequences, producing more diverse data which can be more beneficial for model training. After filtering, the remaining 3,758 AMPs were included in the LMPred dataset, being 24% more than the next largest collection used by Bhadra et al. (1).

There is little incentive to experimentally prove a peptide is non-AMP, and thus there are no large repositories of peptides that have been shown to lack desirable activities. Therefore, without access to a formal non-AMP database, negative samples were collected similarly to prior research (4, 20). Sequences that had been reviewed and verified were downloaded from the UniProt database (https://www.uniprot.org/). The data was then filtered according to the following criteria:

- Any duplicate entries were removed.
- Only samples whose subcellular location was cited as “cytoplasm” were retained to ensure the origination was similar to the AMP samples.
- Any samples labelled as specifically having the activities: “antimicrobial”, “antibiotic”, “antiviral”, “antifungal”, “effector” or “excreted”, were omitted.
- Datapoints with fewer than 10 amino acids, and more than 255 amino acids were removed. The remaining samples then mirrored the range of AMP sequence lengths in the positive sample dataset.
- Sequences which contained unnatural amino acids (Z, B, J, O, U or X) were removed.
- Finally, sequences were screened for similarity using the CD-HIT program, using a 40% similarity threshold. The threshold can be stricter for non-AMPs given the larger number of available sequences, and this ensures a more diverse dataset overall.

Previous research found that if the negative sample sequence length distribution matched that of the positive samples, this resulted in the highest classification accuracy models (20). Therefore 3,758 non-AMPs were selected from the remaining pool of 33,722 to compile a final dataset which matched this criterion. The dataset was then split using sklearn’s “train_test_split” function (https://scikit-learn.org/); 40% of the data was set aside for training, 20% for validation and the remaining 40% for testing. The splitting was performed on a stratified basis and the distribution of samples across these partitions can be found in the supplementary information. The dataset has been made freely available to download at: https://github.com/williamdee1/LMPred_AMP_Dataset.

### Language Model Embeddings

Instructions were followed on the ProtTrans Github page for how to create word embeddings using each LM. An overview of this process is as follows:

- Download the specific tokenizer and pre-trained LM hosted on the ProtTrans Rostlab server.
- Convert the sequences of amino acids into a list and add spaces in-between each amino acid.
- Any unnatural amino acids (“U, Z, O, B, J”) are mapped to “X”.
- The sequence IDs are encoded in batches by the tokenizer, padding with zeros to a specified max length so all inputs are the same length.
- Torch tensors representing the input IDs and the mask used for the attention mechanism are created.
- The embeddings are generated in batches of ten to ensure memory constraints are not breached and the output is saved as a numpy array.

Certain LMs produce special tokens, such as [CLS] or [SEP] tokens, that are included in the embedding array. The CLS token is created by some models to be used as an intelligent average 1-dimensional vector summarizing the full 2-dimensional embedding. It is often used as the input for NLP classification tasks. The SEP token separates any special tokens from the embeddings. In this project the full embeddings were used as not all LMs produce CLS tokens and this ensures greater comparability across results. Additionally, using the full embeddings ensures valuable information isn’t lost and, instead, the neural network can decide how to screen the data through use of the filters, kernel and max pooling layers.

An example of the embedding output is shown in Figure 1.

**Fig. 1.**
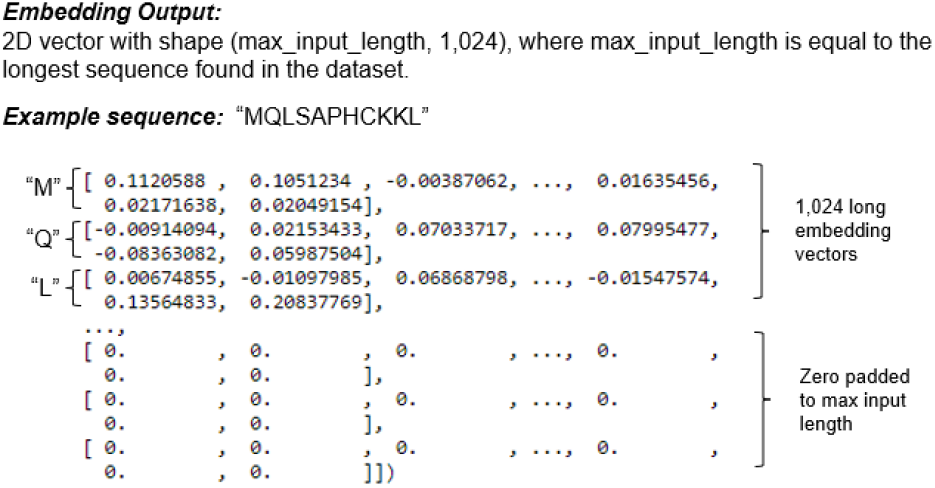
An example of how the sequence ‘MQLSAPHCKKL’ would be represented after applying the pre-trained language models to create an embedding vector.

### Model Architecture

A convolutional neural network (CNN) was chosen as the classifier for this paper, and was built using the Keras framework (https://keras.io/), which utilizes the Tensorflow (37) back-end.

### Embedding Output Example

Two different architectures were tested for each LM. Since the models are used to produce word embeddings there is no need for an initial embedding layer in the CNN, however the number of convolutional, max pooling, dense and batch normalization layers were altered to investigate the impact on performance. A more basic configuration with one convolutional, one max pooling, one batch normalization and then a dense sigmoidal output layer was found to produce the highest classification accuracy model for the BERT UniRef100, BERT BFD and T5 BFD embeddings. A more complex architecture employing two convolutional, two max pooling, two batch normalization, two dense and two dropout layers before the final dense sigmoid output layer, was found to work best for the T5 UniRef50 and XLNet LM embeddings. An illustration of these architectures can be found in Figure 2.

**Fig. 2.**
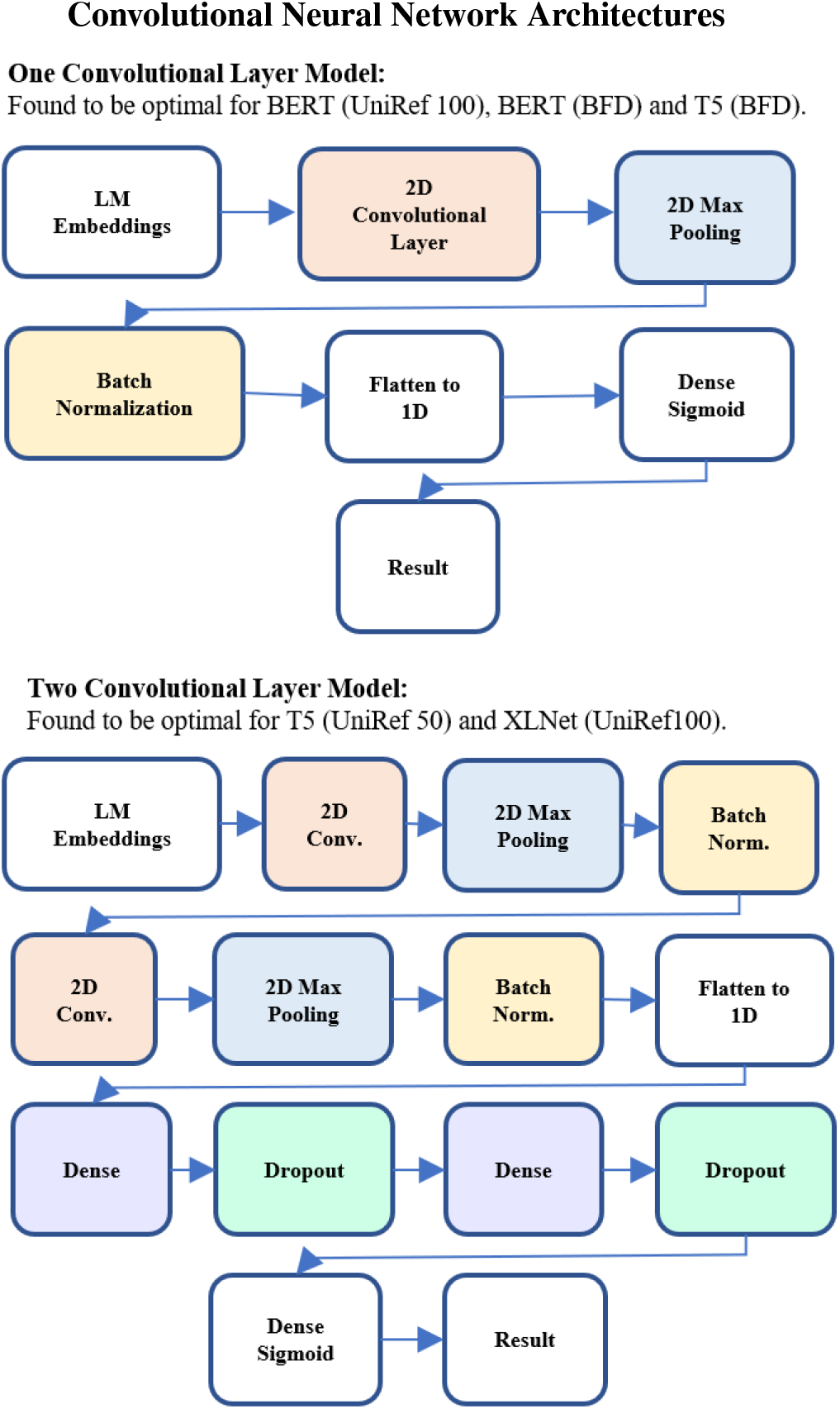
An illustration of the two CNN architectures used with the different language model embedding inputs.

### Hyperparameter Tuning

For each set of LM embeddings, tuning was performed independently to ascertain the optimal hyperparameters for each resulting CNN model. The tuning was performed using the Keras Tuner (https://keras.io/keras_tuner) which provides a framework to apply different search algorithms. Both the Hyperband (38) and the Bayesian Optimization (39) tuning algorithms were used. Utilizing the two methods in combination has been shown to outperform using them separately as, “bandit-based approaches” (like Hyperband) “lack guidance”, whereas Bayesian optimization across the entire search space can be “computationally infeasible” (40). The tuning uses only the training and validation partitions of the data.

Tuning was performed for a maximum of 10 epochs for Hyperband, and 15 for Bayesian Optimization, with an early stopping callback after 6 epochs without a decrease in validation loss, and a reduction of the learning rate by a factor of 1e^-1^ after 4 epochs without a decrease. The optimal parameters were the ones which produced the model with the lowest validation loss during training. The entire search space is detailed in Table 1.

**Table 1.** Hyperparameter Search Space

| Layer | Hyperparameter | Values |
| --- | --- | --- |
| Convolutional | Filter | Min = 32, Max = 512, Step = 32 |
| Convolutional | Kernel | Min = 3, Max = 33, Step = 2 |
| Convolutional | Weight initializer | ‘RandomNormal’, ‘HeNormal’ |
| Convolutional | Activation Function | ‘ReLU’ |
| Max. pooling | Pool size | Min = 2, Max = 14, Step = 2 |
| Max. pooling | Pool strides | Min = 256, Max = 1024, Step = 128 |
| First Dense | Units | Min = 256, Max = 1024, Step = 128 |
| Second Dense | Units | Min = 32, Max = 256, Step = 32 |
| Dropout | Dropout Rate | Min = 0.0, Max = 0.5, Step = 0.1 |
| Optimizer | Learning Rate | Min = $1e^{-7}$, Max = $1e^{-3}$, Step = $1e^{-1}$ |
| Optimizer | Optimizer | ‘Adam’, ‘SGD’ |

### Model training and evaluation

Once hyperparameters were selected for each LM-embedding approach, each CNN model was trained for 30 epochs with the optimizer loss function set to “binary_crossentropy” and metrics set to “accuracy”. Early stopping was set to 12 epochs, and the learning rate was reduced by a factor of 1e^-1^ after 4 epochs without an improvement in validation loss. The model checkpoint callback saved the model with the lowest validation loss during training.

In terms of evaluation, each model was tested for accuracy (Ac.) on the test set for both the Veltri and the LMPred datasets. Accuracy is considered the goal metric as identifying positive and negative samples correctly is equally important. Additionally, sensitivity (Sn.), specificity (Sp.), Matthew’s correlation coefficient (MCC) and the area under the ROC curve (AUC) were also calculated to provide a full overview of model performance. These metrics were calculated as follows using the number of true positive (TN), true negative (TN), false positive (FP) and false negative (FN) predictions:

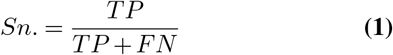

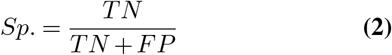

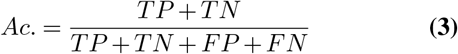

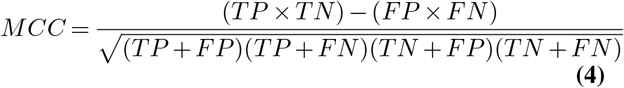

The area under the ROC curve (AUC) was calculated using the sklearn metrics package.

The results were compared to that of the available webserver models, produced by previous state-of-the-art AMP prediction papers, when provided the same test sets. These include; the CAMP server’s artificial neural network, support vector machine, discriminant analysis and random forest models (12), AmPEP’s random forest (1), iAMP-2L’s fuzzy K-nearest neighbour (4), iAMPpred’s support vector machine (17) and Veltri’s CNN model (20).

### Replicating the state-of-the-art approach

In order to ensure the most robust comparison with the current state-of-the-art method for AMP prediction, a replica of the CNN built by Veltri et al. (20) was constructed. Firstly, it was tested on the Veltri dataset, and its efficacy was compared to that of the original paper, to verify it had been replicated correctly. This approach was then tested on the LMPred test set, having been trained using the additional training and validation samples provided by that dataset, which the Veltri webserver model has not had access to. Lastly, since some positive samples in the LMPred dataset were sourced from the dataset used by (20), the webserver model will have been trained on 457 of the samples it is then asked to predict. Metrics will therefore also be included showing the efficacy of the Veltri webserver model at predicting only unseen data.

### Technical settings

Creating word embeddings, as well as hyperparameter tuning, training and evaluation of the CNN models was performed using Google Colab Pro (https://research.google.com/colaboratory/). A Tesla P100-PCIE-16GB GPU was used, as well as up to 24GB of RAM and 150GB of disk resources.

Models using two convolutional layers (T5 UniRef50 and XLNet) took 107 seconds per epoch to train and 166 seconds to load and predict the 3,007 test samples in the LMPred dataset. The size of those models was approximately 1.16GB. This is compared to 35 seconds per training epoch, 34 seconds for testing and 5.2MB in size, for the models using one convolutional layer (BERT UniRef100, BERT BFD and T5 UniRef100).

## Results

### Model performance

#### Veltri Dataset

Table 2 shows the performance of the CNN models, using the different LM embeddings as inputs, when tested on the Veltri dataset. These results have been compared with the results published in (20)’s AMP peptide prediction paper, including the metrics from that paper’s own method, as well as the various webserver models available at that time.

**Table 2.** Comparison with state-of-the-art methods - veltri dataset

| Approach | Method | Sn. (%) | Sp. (%) | Ac. (%) | MCC | AUC (%) |
| --- | --- | --- | --- | --- | --- | --- |
| *Models results from the Veltri et al. (20) paper:* |  |  |  |  |  |  |
| AntiBP2 | SVM | 87.91 | 90.80 | 89.37 | 0.7876 | 89.36 |
| CAMP | ANN | 82.98 | 85.09 | 84.04 | 0.6809 | 84.06 |
| CAMP | DA | 87.08 | 80.76 | 83.92 | 0.6797 | 89.97 |
| CAMP | RF | **92.70** | 82.44 | 87.57 | 0.7554 | 93.63 |
| CAMP | SVM | 88.90 | 79.92 | 84.41 | 0.6910 | 90.63 |
| iAMP-2L | FKNN | 83.99 | 85.86 | 84.90 | 0.6983 | 84.90 |
| IAMPred | SVM | 89.33 | 87.22 | 88.27 | 0.7656 | 94.44 |
| gkmSVM | SVM | 88.34 | 90.59 | 89.46 | 0.7895 | 94.98 |
| Veltri | CNN | 89.89 | 92.13 | 91.01 | 0.8204 | 96.48 |
| *Models created by this paper:* |  |  |  |  |  |  |
| Veltri (Replica) | CNN | 88.48 | 92.70 | 90.60 | 0.8125 | 96.12 |
| BERT (Uni 100) | CNN | 90.31 | 93.82 | 92.06 | 0.8418 | 97.29 |
| BERT (BFD) | CNN | 91.99 | 92.13 | 92.06 | 0.8413 | 97.55 |
| T5 (Uni 50) | CNN | 92.28 | 94.38 | **93.33** | **0.8668** | **97.89** |
| T5 (BFD) | CNN | 88.62 | **95.79** | 92.21 | 0.8463 | 97.41 |
| XLNet (Uni 100) | CNN | 88.48 | 91.01 | 89.75 | 0.7952 | 95.78 |

*Note:* The performance of the webserver models and the Veltri et al. model was sourced from the original Veltri et al. (20) paper. This has been compared with the models built in this paper - being the replica of the Veltri model as well as the models built using different language model embeddings.

The best performing model – T5 pre-trained on the UniRef50 database – achieved an accuracy of 93.33%, being 2.55% higher than that the previous state-of-the-art accuracy results of 91.01% on the same dataset. The T5 UniRef50 model also achieved state-of-the-art MCC and auROC metrics, only being based on specificity by the T5 BFD model, and on sensitivity by the CAMP Random Forest webserver model (12). The auto-regressive model, XLNet, underperformed the two auto-encoder (BERT and T5) LM-based approaches, as well as the approach used by veltri et.al. (20). This underperformance was noted across every metric. However, all models using NLP techniques to create embeddings, and convolutional neural networks as the classifier, displayed higher accuracies than those using biological information and machine learning models.

#### LMPred Dataset

Table 3 shows the performance of the CNN models when tested on the LMPred dataset, compared to the available webserver model predictions when provided the same test data. Similar results were found to Section 3.1.1, with the T5 auto-encoder language model, pre-trained on UniRef50, producing embeddings that resulted in the CNN with the highest accuracy (88.26%).

**Table 3.** Comparison with state-of-the-art methods - LMPred dataset

| Approach | Method | Sn. (%) | Sp.(%) | Ac.(%) | MCC | AUC (%) |
| --- | --- | --- | --- | --- | --- | --- |
| **External Webserver Models:** |  |  |  |  |  |  |
| AmPEP | RF | 56.62 | 43.48 | 50.05 | 0.1050 | n/a* |
| CAMP | ANN | 77.51 | 72.54 | 75.02 | 0.5011 | n/a* |
| CAMP | DA | 81.04 | 71.41 | 76.22 | 0.5269 | 80.16 |
| CAMP | RF | 84.70 | 72.74 | 78.72 | 0.5785 | 83.05 |
| CAMP | SVM | 83.03 | 71.88 | 77.45 | 0.5525 | 80.27 |
| iAMP-2L | FKNN | 76.18 | 74.60 | 75.39 | 0.5079 | n/a* |
| IAMPred | SVM | 89.55 | 55.92 | 72.73 | 0.4828 | 82.28 |
| Veltri | CNN | **90.09** | 66.95 | 78.52 | 0.5863 | 86.45 |
| Veltri (Unseen**) | CNN | 86.42 | 66.95 | 74.94 | 0.5277 | 84.42 |
| **Models created by this paper:** |  |  |  |  |  |  |
| Veltri (Replica) | CNN | 75.98 | 85.11 | 80.55 | 0.6134 | 85.82 |
| BERT (Uni 100) | CNN | 84.43 | 84.91 | 84.67 | 0.6934 | 90.84 |
| BERT (BFD) | CNN | 86.83 | 88.16 | 87.50 | 0.7500 | 93.58 |
| T5 (Uni 50) | CNN | 88.89 | 87.63 | **88.26** | **0.7653** | **94.66** |
| T5 (BFD) | CNN | 86.16 | **88.96** | 87.56 | 0.7515 | 93.68 |
| XLNet (Uni 100) | CNN | 82.63 | 78.92 | 80.78 | 0.6160 | 88.87 |

Note: Results produced when testing models created within this paper, as well as external webserver models, on the LMPred test dataset.
\*n/a: The web server did not output prediction probabilities, so AUC could not be calculated.
\*\*Unseen: Refers to only the samples not already seen by the Veltri webserver model being submitted as the part of the LMPred test set.

The performance difference between the models created by this paper and the state-of-the-art webserver models was more noticeable, likely due to the increased number of samples (111% more than the Veltri dataset) leading to a higher difficulty classification task. The T5 UniRef50 CNN produced 12.4% higher accuracy than the Veltri webserver model, which increased to a 17.8% gap when the Veltri web-server was only presented with unseen test data.

To differentiate between the impact of the increased number of training and validation samples, and the embedding approach taken, a replica of Veltri’s approach was built, utilizing the training and validation data of the LMPred dataset. This approach resulted in an uplift of 2.6% versus the Veltri webserver model, but still fell 9.6% short of the best LM-based CNN’s accuracy - implying that this is the true performance difference between approaches for this dataset.

Consistent with the results found for the Veltri dataset, all models leveraging NLP techniques and CNN’s scored higher accuracies than those utilizing specialist biological feature sets and traditional machine learning models. The T5 model trained on the BFD also displayed the highest specificity, whilst the Veltri and the IAMPred (17) webserver models produced the highest sensitivity. The XLNet embeddings proved to be inferior to both BERT and T5 and did not perform significantly better than the replica Veltri model.

## Discussion

This paper has proposed a novel method for producing model inputs for classifying AMPs. By utilizing the contextualized embeddings produced by pre-trained language models, the resulting convolutional neural network achieved improved classification accuracy compared to the existing state-of-the-art methods across two datasets. This provides further evidence that NLP techniques can replicate some of the “language of life” which previously required extensive time and biological knowledge in the feature engineering stages to represent.

The results support the research of Elnaggar et al. (33) who found that true bi-directionality of contextual embedding was extremely important for protein structure prediction. This implies that the structure of amino acids is important bi-directionally and it is likely that the permutation modelling that XLNet uses, whilst it can work for sentences, is not so applicable in the case of peptide or protein sequences. The more distant dependency modelling that XLNet allows appears to be less beneficial in this use case, as the BERT-based embeddings do not suffer from lacking it.

Furthermore, this research also supported Elnaggar et al. (33)’s results that found T5 to be the superior model for prediction – as the T5 model trained on UniRef50 generated the highest accuracy metric when tested on both datasets. This demonstrates the benefits of using the whole transformer architecture to build the pre-trained language model, rather than just the encoder (BERT) or decoder (XLNet).

Also similar to Elnaggar et al. (33)’s findings, there was no conclusive evidence that pre-training the LMs on larger datasets resulted in embeddings that improved predictive accuracy. On both the Veltri and the LMPred dataset, the T5 model trained on the BFD produced lower accuracy than the one trained on the UniRef50 dataset. The BERT BFD model did outperform the BERT UniRef100 model, but only on the larger, LMPred dataset. More diverse pre-training datasets may prove to be the optimal approach for this problem. This is implied by the results, as UniRef50 only includes the sequences from UniProt that do not share more than 50% sequence similarity. Additionally, the T5 model is also trained on a large, diverse “cleaned” corpus, compared to the smaller and more uniform corpora used to initially produce BERT. The abundance of noise in these databases, i.e., in UniRef100 caused by duplicated sequences, may have proved detrimental to overall learning.

Future research could investigate the effectiveness of the pre-trained language models not considered in scope by this paper. These include the ELECTRA and ALBERT models, as well as the Transformer-XL model when it is released on the ProtTrans Github page (https://github.com/agemagician/ProtTrans). This work would likely provide further support to the evidence found in this paper that auto-encoder models produce more context-rich embeddings which can be more effectively used as inputs into CNN models predicting AMPs. It would be interesting to note whether these performance trends are similar across AMPs with differing activities (i.e. anticancer compared to antihypertensive peptides).

Further work could also split the AMPs into explicit cohorts, i.e., based on sequence length, or the frequency of specific amino acids. This research may reveal that different language models excel at predicting different cohorts, a hypothesis that may be supported by the evidence that longer-term dependency modelling is more important in the case of longer sentences for NLP tasks, or in this case - longer sequences.

## Acknowledgements

I would like to thank Ahmad Abu-Khazneh for his feedback and guidance during the completion of this project.

## Supplementary Information

### 1. Sequence Length Distributions – LMPred Dataset

**Figure S1:**
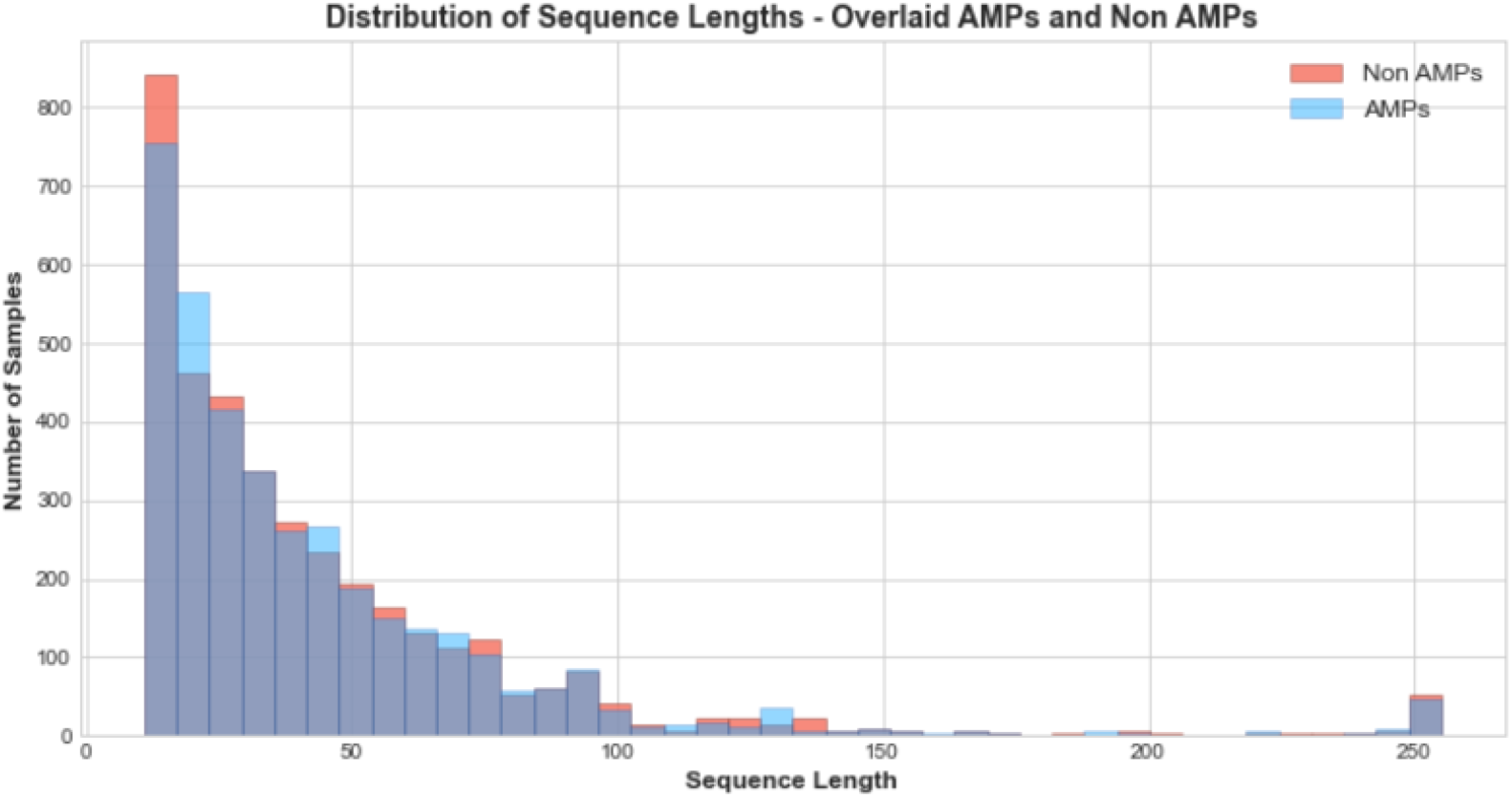
Sequence length distributions of the 3,758 AMP (blue) and 3,758 non-AMP (red) samples in the full LMPred dataset.

**Figure S2:**
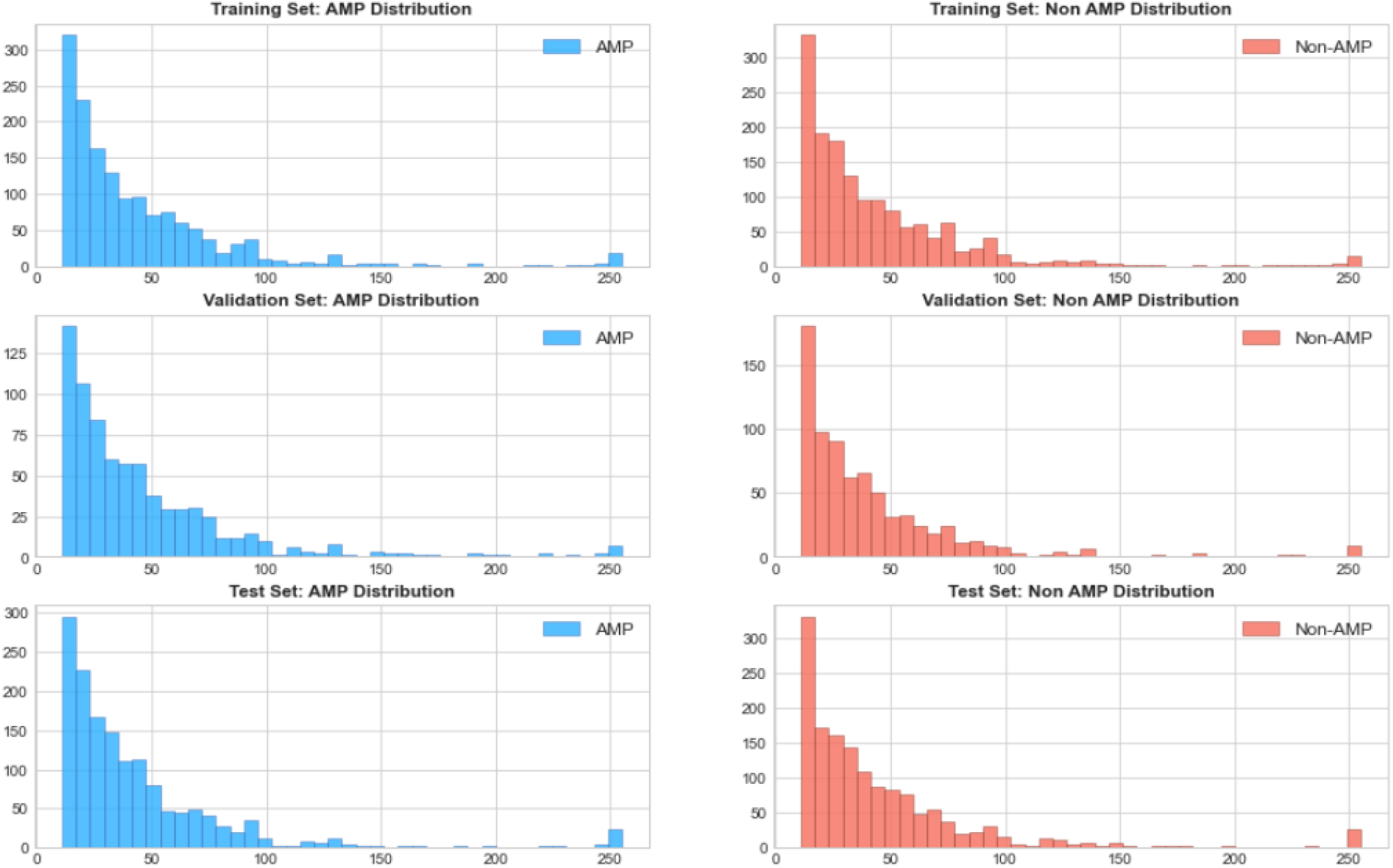
Sequence length distributions of the AMPs and non-AMPs across the training (40%), validation (40%) and test (20%) datasets.

### 2. Mean Embedding Values

Below shows some evidence of the different information conveyed by the embeddings produced by the different pre-trained language models. This is most obvious when looking at the embedding of “L” in position 11 of the non-AMP sample, compared to it’s embedding in position 10 of the AMP sample. This difference is most stark for the T5 models where “L” is given the average value of -0.002 in the non-AMP and -0.010 in the AMP position. Overall, the T5 models appear to display the most absolute variance between embeddings.

**Figure S3:**
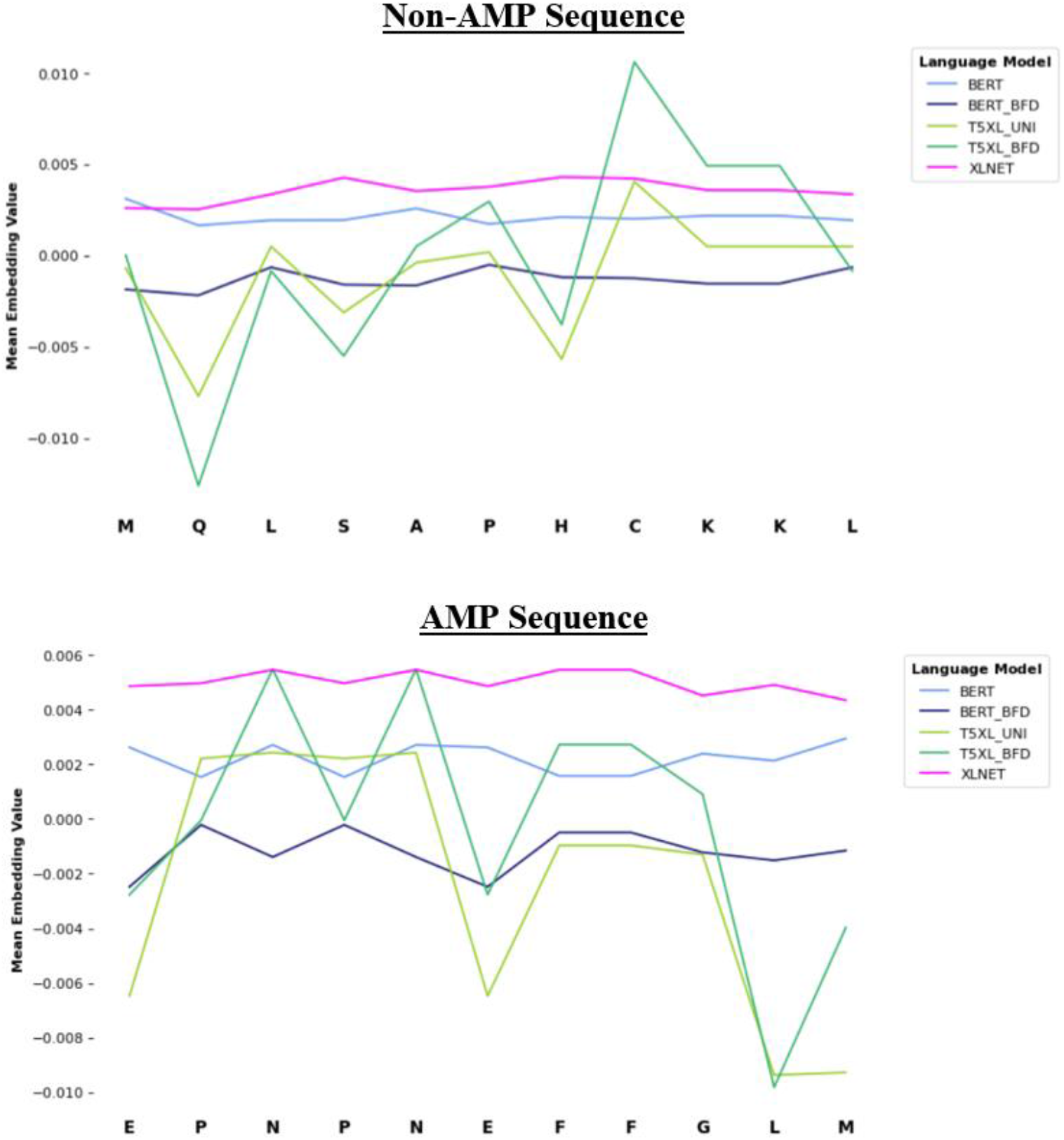
Line plots showing the comparative average value per amino acid assigned by the different pre-trained language models to an AMP and non-AMP 11 amino acids in length.

### 2. Data Availability

The LMPred dataset, along with notebooks detailing how the research can be replicated, can be found at the following Github page: https://github.com/williamdee1/LMPred_AMP_Dataset

